# Sounds are remapped across saccades

**DOI:** 10.1101/839449

**Authors:** Martin Szinte, David Aagten-Murphy, Donatas Jonikaitis, Luca Wollenberg, Heiner Deubel

## Abstract

To achieve visual space constancy, our brain remaps eye-centered projections of visual objects across saccades. Here, we measured saccade trajectory curvature following the presentation of visual, auditory, and audiovisual distractors in a double-step saccade task to investigate if this stability mechanism also accounts for localized sounds. We found that saccade trajectories systematically curved away from the position at which either a light or a sound was presented, suggesting that both modalities are represented in eye-centered oculomotor centers. Importantly, the same effect was observed when the distractor preceded the execution of the first saccade. These results suggest that oculomotor centers keep track of visual, auditory and audiovisual objects by remapping their eye-centered representations across saccades. Furthermore, they argue for the existence of a supra-modal map which keeps track of multi-sensory object locations across our movements to create an impression of space constancy.

## Introduction

The location of sounds cannot be directly determined from the auditory input received at the ears, but must instead be derived from a series of complex computational steps (King, Schnupp, & Doubell, 2001). Indeed, sounds are localized from the integration of acoustical (inter-aural time difference and interaural level difference) and spectral cues within a network of midbrain (King, 2004; Sparks & Nelson, 1987) and cortical structures (Arnott & Alain, 2011), resulting in head-centered encoding of auditory space. However, it has been shown that the representation of sounds gradually shifts from a head-centered to an eye-centered reference frame between the inferior colliculus —IC— and the superior colliculus –SC—(Bulkin & Groh, 2012; Groh, Trause, Underhill, Clark, & Inati, 2001; Jay & Sparks, 1984; 1987; King, Hutchings, Moore, & Blakemore, 1988; J. Lee & Groh, 2012). Furthermore, sound location is also encoded in eye-centered coordinates within several other cortical structures (Grunewald, Linden, & Andersen, 1999; e.g. Lateral Intraparietal area —LIP–: Linden, Grunewald, & Andersen, 1999; or the Frontal Eye Fields —FEF—: Russo & Bruce, 1994; Stricanne, Andersen, & Mazzoni, 1996).

Such a transformation suggests a role for eye-centered representation in auditory localization. In line with this idea, Doyle and Walker (2002) showed that, even when instructed to saccade to visual targets and ignore auditory distractors, saccade trajectories still curve away from the location of auditory distractors presented before movement initiation. Accordingly, to induce this systematic effect on the saccade trajectory, the location of the auditory distractor must be represented within the eye-centered oculomotor centers used for saccade targeting. It is believed that the simultaneous representation of the auditory distractor and the visual saccade target locations within the same oculomotor map leads to competition during the elaboration of the motor plan (Kruijne, Van der Stigchel, & Meeter, 2014).

This competition then results in systematic changes in the curvature of the saccade trajectory as a function of the distractor location. However, if auditory distractors are indeed represented within eye-centered oculomotor centers, then what would happen when the mapping between head-centered and eye-centered encoding is changed because the eyes have moved but the head (and ears) have remained stable?

When the eyes move, even an object that is stationary in the external world will project onto a different part of the retina after the movement. For visual objects, different studies have shown that both cortical and sub-cortical structures remap (or update) their visual neuron receptive fields before the movement of the eyes begins (Duhamel, Colby, & Goldberg, 1992; Sommer & Wurtz, 2006; M. F. Walker, Fitzgibbon, & Goldberg, 1995). Visual neurons, which after a saccade will receive an object within their receptive field, show a predictive remapping activity that allows them to keep track of visual objects across eye movements (Wurtz, 2008). Until now, however, remapping effects have only been reported for visually defined objects.

Nevertheless, auditory cells within the SC are modulated by the position of the eyes, with reduced or abolished activity observed when a change in eye direction brings the sound source outside of the eye-centered receptive field of the recorded neuron (Jay & Sparks, 1984; Populin, Tollin, & Yin, 2004). Additionally, it has been shown that humans can accurately shift their gaze toward the origin of an auditory target presented before a movement, even after making a double-step eye or combined eye and head movement (saccade: Goossens & Van Opstal, 1999; or eye-head combined movement: Van Grootel, Van Wanrooij, & Van Opstal, 2011; Vliegen, Van Grootel, & Van Opstal, 2004). The ability to correctly perform such a task suggests that the memorized location of the auditory stimulus was accurately updated, or remapped, across the first saccade such that the second saccade could be precisely targeted. However, these results do not exclude the possibility that the auditory location was represented solely within head-centered auditory maps and simply relayed to eye-centered oculomotor centers (before the onset of the first movement) to form a memorized, eye-centered motor plan. As such, earlier studies leave open the question of the mechanism used to process sound location across saccades.

Here we aim at determining the mechanism used to represent visual, auditory, and audiovisual distractors across saccades. To do so, we built a custom screen (Figure 1A) that allowed us to accurately record eye movements while auditory or visual distractors were presented at different locations. We instructed participants (n = 8) to move their eyes following a double-step sequence consisting of a first (left- or rightwards) saccade to the screen center followed by a second (up- or downwards) saccade along the screen vertical meridian. Distractors were either presented (Figure 1B) before (pre-saccadic distractor trials) or after the first horizontal saccade (inter-saccadic distractor trials). For these two types of trials, we analyzed the second vertical saccade as a function of the distractor position and sensory modality. We predicted that if a distractor was represented within eye-centered oculomotor centers, its representation would compete with the representation of the visual saccade target and influence the trajectory of the second saccade (Kruijne et al., 2014; Sheliga, Riggio, & Rizzolatti, 1994). For inter-saccadic trials, in which screen (or head-centered) and eye-centered representations of the distractor are equivalent, we simply predicted that the saccade trajectory would curve away from the distractor. As expected, in these inter-saccadic distractor trials for which both eye- and head-centered representations of the distractor were aligned, we found that the trajectory of the second saccade systematically curved away from the distractor screen location irrespective of whether the distractor was visual, auditory or audiovisual.

**Figure 1.**
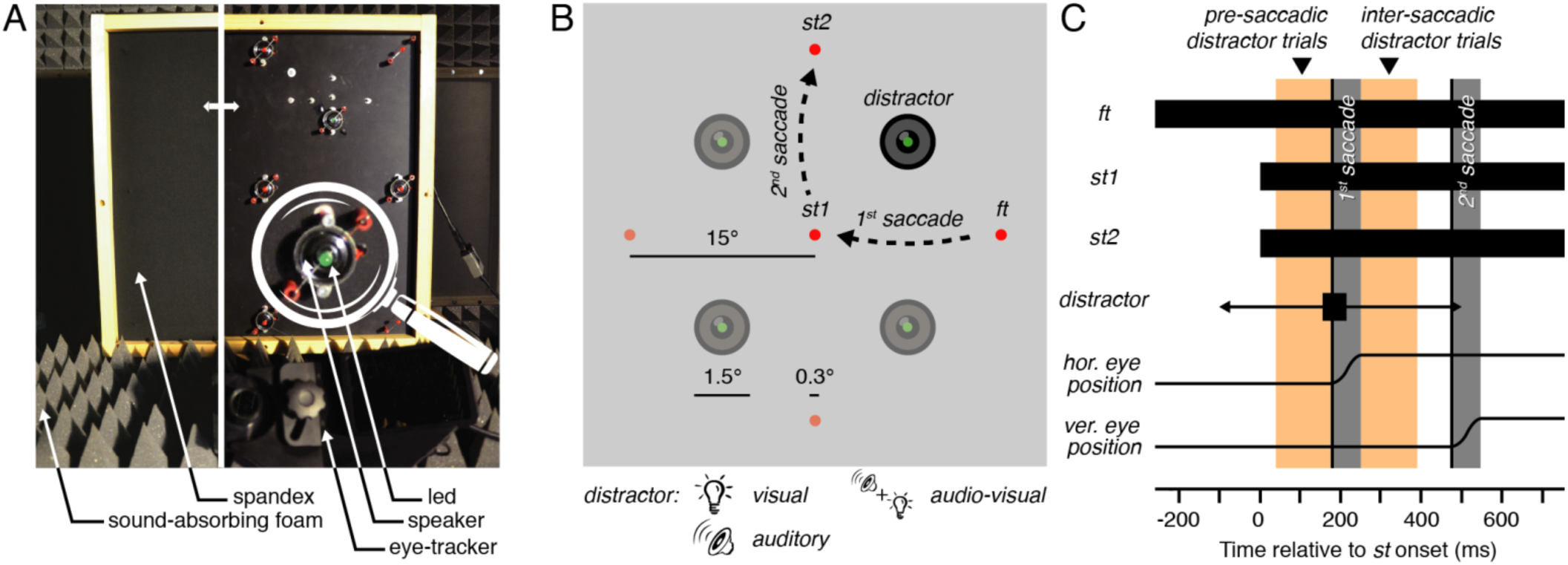
Apparatus and stimulus timing. **A.** We built a combined eye-tracking and audiovisual screen consisting of LEDs and sound speakers (the left side of the panel shows the setup covered with spandex, transparent to sounds and lights, while the right side shows the equipment below) in a cabin covered with sound-absorbing foam. **B.** The presentation of the fixation target (*ft*) was followed by the simultaneous appearance of the saccade targets (*st1* and *st2*). Participants (n = 8) executed a sequence of two saccades while ignoring the onset of a distractor, a brief (50 ms) visual, auditory or audiovisual stimulus presented at a location either clockwise or counterclockwise relative to the second saccade direction. **C.** Pre-saccadic and inter-saccadic trials (orange bars) were defined according to the presentation time of the distractor: the period preceding (pre-saccadic) or following (inter-saccadic) the first saccade. The curvature of the second saccade of pre-saccadic and inter-saccadic distractor trials was analyzed as a function of the distractor position and sensory modality (visual, auditory or audiovisual).

For pre-saccadic trials, for which the distractor occurred before the first saccade, we could formulate two separate hypotheses about the curvature of the second saccade. If the eye-centered (or retinotopic) distractor representation is not being remapped across the first eye movement, we would predict that the second saccade curves *toward* the screen distractor position (because this corresponds to curvature away from the retinal position of the distractor before the first saccade). On the contrary, if the eye-centered distractor representation is being remapped across the first eye movement, we would predict that the second saccade curves *away* from the distractor screen position. Supporting the second hypothesis, we found that vertical saccades recorded in pre-saccadic trials, systematically curved away from the distractor screen location irrespective of its sensory modality. Together, these results show that both auditory and visual objects are registered within eye-centered oculomotor centers. Moreover, they show that when the eye position changes, stimulus representations are remapped, ensuring that eye-centered representations of stimuli from different modalities can all maintain their alignment to the external world.

## Results

We first focused on inter-saccadic distractor trials (Figure 2). In these trials the distractor occurred after the first saccade but before the onset of the second saccade. Analysis of saccade curvature as a function of the distractor position can therefore inform us on whether a brief visual and/or auditory stimulus was represented within eye-centered oculomotor centers (Figure 2A). Figure 2B shows the averaged second saccade trajectory and curvature angle normalized relative to trials without distractors as a function of the distractor sensory modality (see Methods). We found that vertical saccades curved away from the screen position of the visual distractor when it was presented during the inter-saccadic interval. This effect was consistent across trials, as made evident by the comparison between the normalized curvature angle observed for visual inter-saccadic distractor trials (Figure 2B left panels: −2.65 ± 0.41°—mean ± SEM—) and trials without distractor (0.30 ± 0.11°, *p* < 0.0001). Interestingly, auditory distractors produced smaller but similar effects. Saccade trajectories systematically curved away from the screen location where inter-saccadic sounds were played (Figure 2B middle panels: −0.52 ± 0.14°, *p* < 0.0001). Finally, audiovisual inter-saccadic distractors had similar effects on the curvature of the second saccade and resulted in systematic curvature away from the screen position where they were presented (Figure 2B right panels: −3.34 ± 0.53°, *p* < 0.0001). Altogether, these results suggest that auditory and/or visual stimuli are represented within eye-centered oculomotor centers and compete with visual saccade targets, resulting in a systematic bias of saccade trajectory curvature.

**Figure 2.**
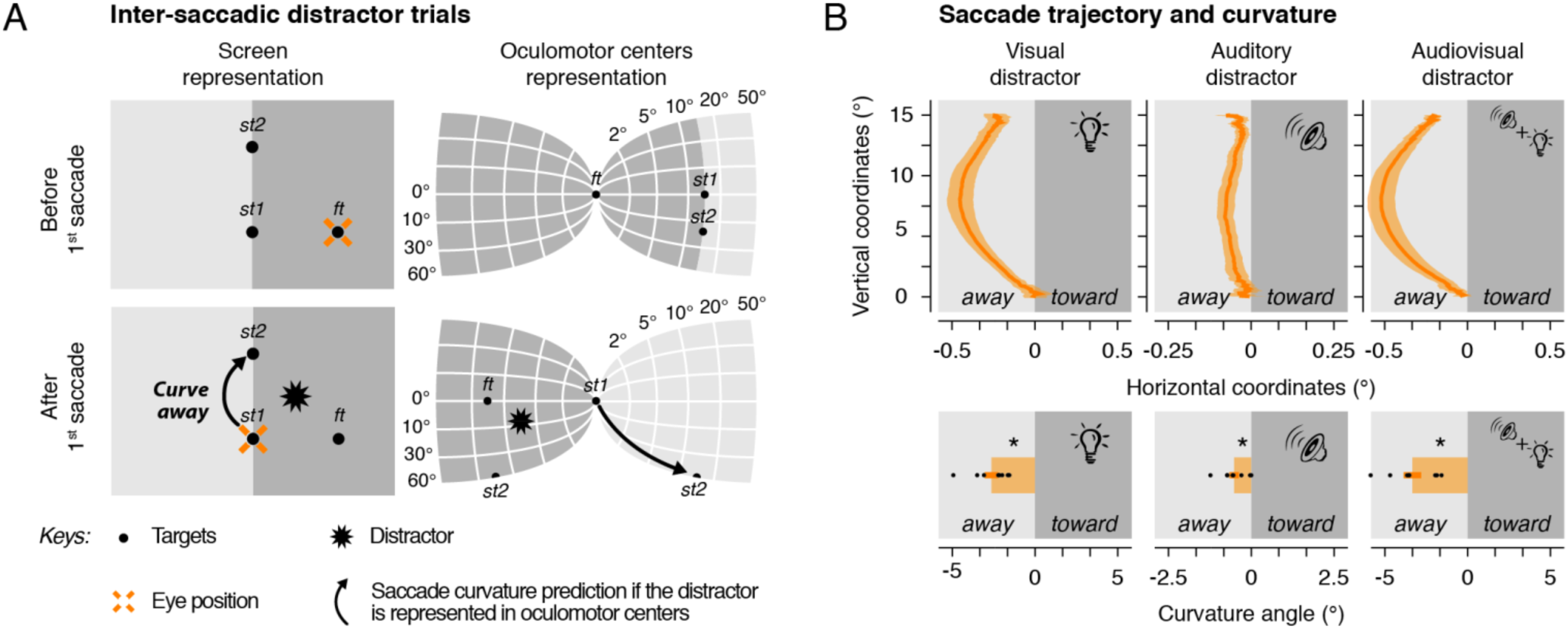
Inter-saccadic distractor trials. **A.** Screen (left panels) and oculomotor centers (right panels) representation of the targets (black dots), the eye position (cross), the distractor (star) and the saccade curvature prediction (arrow) before (top panels) and after (bottom panels) the first horizontal saccade. If the distractor is represented in oculomotor centers, we predicted that the second saccade would curve away from the distractor screen position. **B.** Averaged normalized second saccade trajectory (top panels) and curvature angle (bottom panels) observed following the presentation of a visual (left panels), an auditory (center panels) or an audiovisual (right panels) inter-saccadic distractor. Saccade trajectories are rotated in order to have upward saccades and negative x-values representing coordinates away from the screen position of the distractor. Areas around the averaged saccade trajectory and error bars represent SEM. Black dots show individual participants. Asterisk indicates a significant effect (*p* < 0.05, ns: non-significant). Note that we illustrated two known features of oculomotor centers: cortical magnification factor (see logarithmic scale) and the double visual inversion (left-right and top-bottom) relative to the screen representation.

Next, we focused on trials in which the distractor preceded the two saccades. Depending on the position of the targets and of the distractor on the screen, we labeled these trials as inter-hemifield or intra-hemifield pre-saccadic trials. Inter-hemifield pre-saccadic trials are characterized by distractors horizontally presented at a position laying in between the fixation and the first saccade target (Figure 3A). For these trials the first saccade shifts the distractor’s eye-centered representation in the opposite visual hemifield relative to where it was originally presented (i.e. inter-hemifield). Thus, these trials allow us to determine if oculomotor centers, while processing the second saccade, take the execution of the first saccade into account when representing visual, auditory and audiovisual distractors. If the displacement from the first horizontal saccade is not taken into account, then we would predict that vertical saccades curve toward the distractor screen position. On the contrary, if the movement of the first saccade is considered, such that the distractor representation is being remapped across the first saccade, we would predict that the second saccade curves away from the distractor screen position.

**Figure 3.**
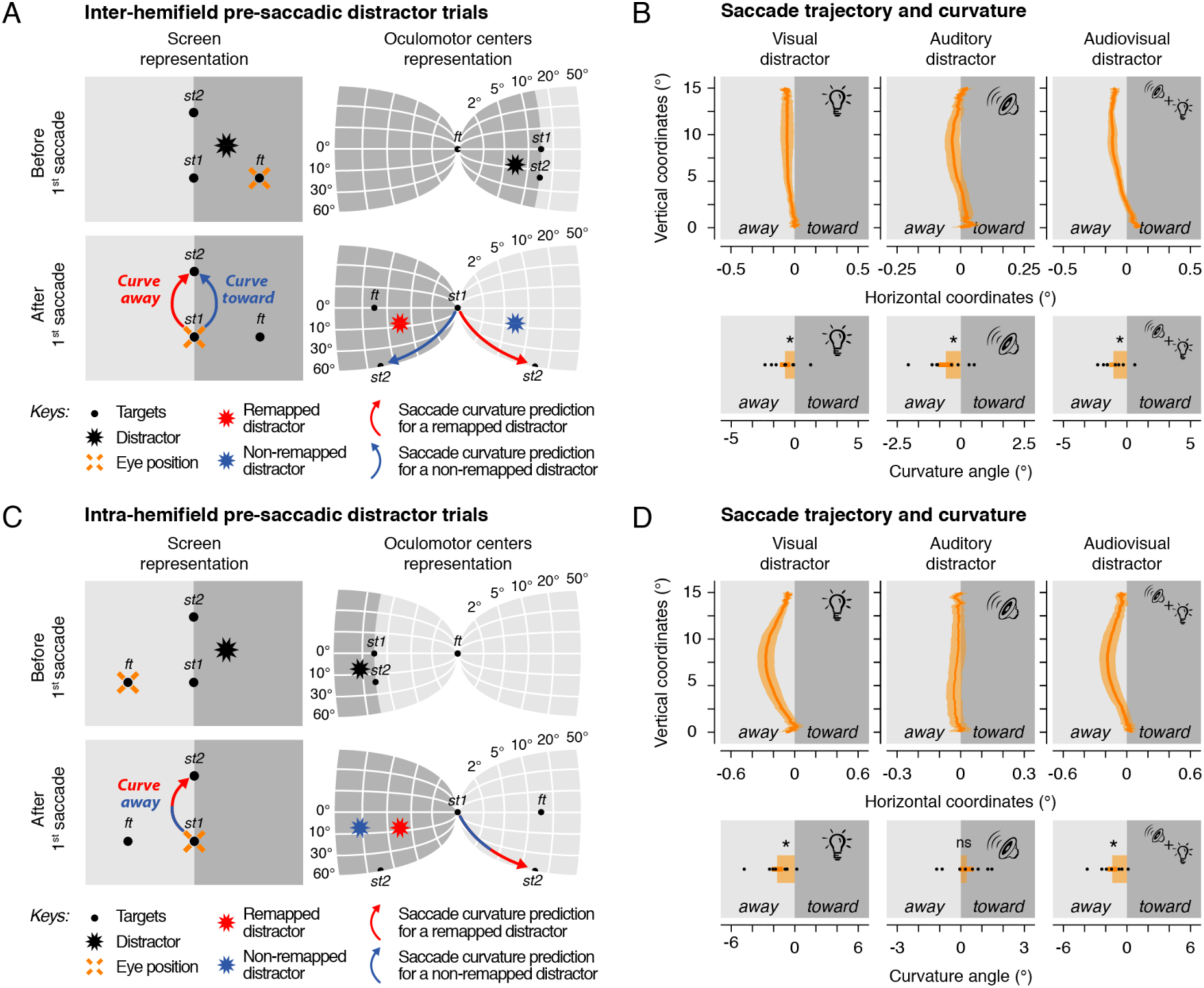
Inter- and intra-hemifield pre-saccadic distractor trials. **A-C.** Screen and oculomotor center representations before and after the first saccade. For inter-hemifield pre-saccadic trials (A), the distractor is presented near to the initial fixation, located in the same hemifield. Here, if oculomotor centers remap the distractor across saccades, we predicted that vertical saccades would curve away from the distractor screen position (red arrow), otherwise they will curve toward it (blue arrow). In intra-hemifield pre-saccadic trials (C), the distractor is presented further from the initial fixation position, located in the other hemifield. We predicted that vertical saccades would curve away from the distractor screen position irrespective of whether oculomotor centers remap (red arrow) the distractor across saccades or not (blue arrow). **B-D.** Averaged normalized second saccade trajectory (top panels) and normalized curvature angle (bottom panels) observed following the presentation of a visual (left panels), an auditory (center panels) or an audiovisual (right panels) inter-hemifield (B) and intra-hemifield (D) pre-saccadic distractor. Conventions are as in Figure 2.

We found that the presentation of inter-hemifield pre-saccadic distractors resulted in saccade curvature away from the distractor screen position. This effect was found both for visual (Figure 3B left panels: −0.76 ± 0.40°, *p* = 0.0130), auditory (Figure 3B middle panels: −0.58 ± 0.30°, *p* < 0.0026) and audiovisual distractors (Figure 3B right panels: −1.05 ± 0.33°, *p* < 0.0001). These results strongly suggest that auditory and visual representations within eye-centered oculomotor centers are remapped across saccades to compensate for the effects of the first horizontal eye movement.

Next, we analyzed intra-hemifield pre-saccadic trials. These trials are characterized by distractors presented at locations laying further away from the fixation and the saccade targets (Figure 3C). For these trials the first horizontal saccade shifts the distractor’s eye-centered representation within the same visual hemifield relative to where it was originally presented (i.e. intra-hemifield). Contrary to inter-hemifield pre-saccadic trials, these trials do not allow us to determine if oculomotor centers consider the movement of the first saccade when processing the second saccade. Indeed, we predicted that vertical saccades would curve away from the distractor screen position irrespective of whether the distractor representation was remapped. We found that the presentation of intra-hemifield pre-saccadic distractors resulted in saccade curvature away from the distractor screen position. This effect was, however, only observed systematically across participants for visual (Figure 3D left panels: −1.38 ± 0.45°, *p* < 0.0001) and audiovisual distractors (Figure 3D right panels: −1.15 ± 0.37°, *p* < 0.0001). For auditory distractors, although the averaged saccade trajectory curved away from the distractor screen position, the curvature angle observed across participants did not significantly differ from trials without distractors (Figure 3D middle panels: 0.24 ± 0.28°, *p* = 0.8308). Interestingly, as sounds were presented at the same screen position in inter- and intra-hemifield pre-saccadic trials, this latter effect suggests that the localized sound representations within oculomotor centers were modulated by the eccentricity between the position of the eyes and the sound source.

Altogether, we found that vertical saccades curved away more when a visual distractor was presented after the first saccade (inter-saccadic trials) rather than before the first saccade (pre-saccadic trials) for both inter-hemifield (*p* < 0.0001) and intra-hemifield trials (*p* = 0.0004). The same effects were found for audiovisual distractors (*ps* < 0.0001) and for auditory intra-hemifield distractors (*p* = 0.0084). However, auditory inter-hemifield pre-saccadic distractor trials did not differ from auditory inter-hemifield inter-saccadic distractor trials (*p* = 0.8206). As pre-saccadic distractors were necessarily presented earlier than inter-saccadic distractors relative to the onset of the second saccade, the reduced influence of pre-saccadic distractors on the second saccade trajectory most likely reflects the effect of time between the distractor presentation and the execution of the movement. Such an influence of time on the strength of the curvature, demonstrated for visual distractors (an effect previously shown for visual distractors: Jonikaitis & Belopolsky, 2014), might be less visible for less well localized objects such as the auditory distractor we used here.

## Discussion

We behaviorally probed the neural representation of visual, auditory and audiovisual objects using objective measures of saccade trajectory. Our custom-made screen and paradigm allowed us to determine the reference frame in which these different stimulus modalities were represented using a double-step saccade task. We found that the curvature of vertical saccades was systematically biased by visual, auditory and audiovisual distractors briefly presented before movement onset. Moreover, the trajectory of the second, vertical saccade was found to curve away from the screen position of the distractor when the execution of the first saccade shifted its eye-centered representation in the opposite visual hemifield. This systematic influence of the distractors on the saccade trajectory curvature suggests that both auditory and visual distractor representations competed with the visual saccade targets in eye-centered oculomotor centers. Importantly, our results indicate that oculomotor centers keep track of both sound and light positions by remapping their eye-centered representations across saccades.

Saccade trajectory has previously been shown to reflect not only individual idiosyncrasies, but also the spatial layout of the scene over which the eyes move (Van der Stigchel, Meeter, & Theeuwes, 2006). Here, we were able to reveal changes in the saccade trajectory as a function of the distractor interval, position, and sensory modality. Previous studies indicated that the presence of a salient visual or auditory distractor influences saccade trajectories by causing systematic deviations and curvature (Doyle & Walker, 2002; Frens & Van Opstal, 1995; Heeman, Nijboer, Van der Stoep, Theeuwes, & Van der Stigchel, 2016; Sheliga et al., 1994). While curvature away from the distractor reflects an inhibition of the memorized distractor location (Theeuwes, Olivers, & Chizk, 2005), the occurrence of this systematic effect by itself provides evidence for the registration of corresponding stimuli within oculomotor centers (Kruijne et al., 2014). As oculomotor centers responsible for saccade planning and execution are known to be organized in an eye-centered reference frame (Robinson, 1972; Robinson & Fuchs, 1969), our results suggest that both, visual and auditory distractor stimuli were processed by eye-centered multi-sensory neurons within these areas.

Such multimodal neurons have been documented in awake macaques (Jay & Sparks, 1984; 1987), especially within the deeper layers of the SC, a site where both modalities have been found to converge to form a common motor map (Frens, Van Opstal, & Van Der Willigen, 1995; King, 2004). Interestingly, the response of collicular neurons depends both on the position of the sound source and on the position of the eyes in the orbit (Jay & Sparks, 1984; 1987). In other words, although sounds are initially encoded in a head-centered reference frame (King, 2004; Sparks & Nelson, 1987), collicular neurons respond differently as a function of the gaze direction, suggesting that the existence of eye-centered auditory receptive fields (Jay & Sparks, 1984; 1987). These converging neurophysiological studies provide further evidence for the encoding of multimodal stimuli within eye-centered oculomotor centers.

We propose that the collicular or cortical eye-centered registration of the auditory and visual distractors, and the remapping of their eye-centered representations across the saccade could explain the effects observed here. Indeed, if sounds are represented within a common, eye-centered reference frame, their neural representations should be remapped across saccades. Accordingly, when programming the movement of the second saccade, the competition occurs between the saccade target and the remapped distractor location. Thus, because the shift in the eye position due to the saccade is considered, the trajectory of the saccade will deviate away from the screen (i.e. external) location of the distractor.

Such neural remapping of stimulus-related activity has been observed for visual neurons in several different oculomotor centers (Duhamel et al., 1992; Sommer & Wurtz, 2006; M. F. Walker et al., 1995). Remapping consists of a predictive increase in the spiking activity of neurons, which, after the completion of a given eye movement, receive either a salient object or its memory trace in their receptive field (Gottlieb, Kusunoki, & Goldberg, 1998). Moreover, the activity of neurons, which will no longer receive the salient object in their receptive field after completion of the saccade, has been found to predictively return to a baseline level (Colby, Duhamel, & Goldberg, 1996; Duhamel et al., 1992; Kusunoki & Goldberg, 2003; Nakamura & Colby, 2002). However, no study published to date has reported such a neural remapping of auditory objects occurring across saccades.

In a series of experiments recording single cells in the superior colliculus of awake macaques, Jay and Sparks (Jay & Sparks, 1984; 1987) demonstrated that eye-centered auditory receptive fields, when activated by a specific external sound source, reduce their firing after a change in eye coordinates. While these studies demonstrate that the activity of auditory neurons is modulated by the gaze direction, the authors did not record activity before a saccade and thus could not demonstrate a potential remapping of eye-centered auditory neurons.

In our experiment, we found systematic saccade curvature away from the sound distractor when the first saccade shifted the sound distractor into the opposite visual hemifield (inter-hemifield). We argue that these results are compatible with the view that sounds are encoded at least partially within eye-centered oculomotor centers and remapped across the first saccade. This effect was, however, less systematic when the first saccade shifted the sound distractor within the same visual hemifield (intra-hemifield). One crucial difference between these two conditions is the eccentricity between the fixation target and the distractor position before the first saccade. During intra-hemifield trials the distractor occurred on the opposite side of the display relative to the current gaze position. Thus, the eccentricity of the distractor from the eye was about twice as large in intra-hemifield trials (distance between fixation target and distractor: ~21°) relative to inter-hemifield trials (~11°). Nevertheless, distractors of both trial types were presented at the same screen position (and as such the sounds were equally eccentric relative to the position of the ears). As only the eye-centered eccentricity differed, we argue that the difference we observed here shows that sounds are represented in eye-centered coordinates.

Interestingly, when measuring collicular receptive field shifts as a function of the eye direction, Jay and Sparks found that the spatial shift was less precise for auditory neurons than for visual neurons (Jay & Sparks, 1984; 1987). This effect could be due to a difference in saliency between the stimuli (Alais & Burr, 2004) or due to an intrinsic difference in localization capacity between visually and auditory defined objects. In our experiment sound sources were separated on the screen by 15°, a distance far above auditory localization thresholds observed for example when participants had to discriminate two sounds (Maddox, Pospisil, Stecker, & Lee, 2014) or to saccade toward them (Goossens & Van Opstal, 1999). Nevertheless, the lower precision with which auditory stimuli are localized relative to visual stimuli would be expected to reduce the strength of competition between the auditory distractor and the visual saccade target. In addition to auditory stimuli being localized less precisely than visual ones, the absolute localization of auditory stimuli is also known to be biased toward the current gaze location (Krüger, Collins, Englitz, & Cavanagh, 2016; Pavani, Husain, & Driver, 2008), with the magnitude of the bias increasing with eccentricity. While providing further support for interactions between auditory representations and gaze direction, this effect may also have resulted in an inwards localization bias. If such bias occurs be the case, in the intra-hemifield pre-saccadic condition, the condition in which the eyes start further away from the distractor, sounds remapped representation would lie along the trajectory of the second saccade, rather than to one side of it. One would predict that, overall, this condition would be associated with an absence of systematic curvature away or toward the distractor screen position, as observed in our experiment.

Although the difference between sensory modalities may reflect different localization accuracy, only the remapping of an eye-centered representation of the sound across the saccade can fully explain the observed effects. It has been shown that humans can accurately execute double-step eye movements or eye and head movements toward memorized auditory targets (Goossens & Van Opstal, 1999; Van Grootel et al., 2011; Vliegen et al., 2004). Contrary to our results, these earlier findings, as well as previous studies measuring the effect of sound on saccade curvature using a single eye movement task (Doyle & Walker, 2002; Heeman et al., 2016), could be explained by a mechanism in which auditory targets are initially processed in head-centered auditory maps before being transferred into an eye-centered oculomotor map at a later point in time. In contrast, our effect was observed in a double-step saccade task with distractors presented during saccade preparation. Further, the significant increase of curvature angle found for audiovisual distractors argues for the existence of a single eye-centered reference frame in which distractors were encoded, regardless of their sensory modality. Future studies combining multiple distractors presented before and after the first saccade could be used to more directly test the existence of a single reference frame and to further examine integration across modalities.

The location-dependent effect of auditory distractor stimuli on the curvature of eye-movements indicates that some representation of auditory stimuli must necessarily be encoded within eye-centered oculomotor maps in order to provide the competition that biases the saccade trajectory. We argue that this eye-centered auditory information is present from the initial encoding of the stimulus, with its eye-centered representation being remapped after the first saccade to account for the displacement of the eyes. However, an alternative account is that the auditory information is represented in head-centered spatiotopic maps (Andersen, Essick, & Siegel, 1985; Melcher & Colby, 2008), with the representation converted into an eye-centered reference frame “on-demand” (Henriques, Klier, Smith, Lowy, & Crawford, 1998; Klier, Wang, & Crawford, 2001; J. Lee & Groh, 2012) immediately prior to the second saccade. Under this alternative hypothesis, the representation of the auditory distractor stimulus would not be stored within eye-centered coordinates and thus not be subject to remapping across the saccade. We believe this alternative is unlikely for two reasons. First, we observed a clear effect of curvature exhibited in response to localized auditory distractors. This clearly demonstrates that ocular information has a role in determining the degree of curvature, which is difficult to explain if the auditory distractors were represented solely in a head-centered reference frame. Second, given that the auditory stimuli represent a task-irrelevant distractor that individuals are explicitly instructed to ignore, it is unclear why they would actively convert head-centered information into an eye-centered oculomotor representation after the first saccade. Instead, if the representations are separate, it would be far more advantageous to not perform this transformation, rendering the saccade unaffected by the irrelevant auditory distractor. In summary, while only the direct recording of oculomotor multi-sensory neurons, for example within the SC, could confirm the existence of neural remapping of sounds in eye-centered maps, the pattern of data we observed with auditory, visual and audiovisual distractors best supports a model in which different modalities are tracked in a single eye-centered visual reference frame (Frens et al., 1995; King, 2004).

Such a supra-modal topological map could allow multi-sensory space constancy via predictive remapping of locations as a function of their ability to attract spatial attention (Cavanagh, Hunt, Afraz, & Rolfs, 2010; Rolfs & Szinte, 2016; Szinte, Jonikaitis, Rangelov, & Deubel, 2018) rather than as a function of their modality. As an alternative, one could propose that multiple reference frames are simultaneously maintained. In this framework, auditory representations would be accounted for by eye-centered areas but would also be kept in a head-centered reference frame within other cortical areas (Collins, Heed, & Röder, 2010). Indeed, we found a stronger bias for inter-saccadic than pre-saccadic distractor trials. As only within inter-saccadic trials both the head- and eye-centered representations are aligned, this effect goes in line with the idea that the two reference frames exist separately and converge in oculomotor centers to affect the saccade path. Such an integration, which might enhance the distractor signal and consequently the curvature bias, was previously described at the electrophysiological (Chen, DeAngelis, & Angelaki, 2013) and the behavioral level (Badde & Heed, 2016) in experiments without saccades or sounds.

Finally, our results show that saccade trajectory curvature is affected differently by the two modalities. Nevertheless, inference based on the comparison of saccade trajectory curvature effects across modalities is somewhat limited as we kept stimulus intensity constant, confounding arousal and sensory modality factors. Future work systematically varying the level of the two modalities would better test multi-sensory integration (Godfroy-Cooper, Sandor, Miller, & Welch, 2015).

In conclusion, we found that localized visual and auditory objects are treated on a supra-modal eye-centered oculomotor map and are being maintained in space across saccades via a remapping mechanism.

## Material and methods

### Participants

Eight students and staff members from the Ludwig-Maximilians-Universität München participated in the experiment (age 24-30, 3 females, 2 authors) for a compensation of 10 Euros per hour of testing. All participants except the authors were naive as to the purpose of the study and all had normal or corrected-to-normal vision and audition. The experiment was undertaken with the understanding and written consent of all participants and carried out in accordance with the Declaration of Helsinki. The protocol was designed according to the ethical requirements specified by the LMU München and the approval of an institutional ethics committee for experiments involving eye tracking. All files are available from the OSF database URL: https://osf.io/h3qdf/.

### Setup

Participants sat in a dim-light, sound-attenuated cabin, gazing toward the screen center with their head positioned and kept steady by a chin and forehead rest. The cabin’s walls, floor, ceiling and all other large objects in the room were covered with Flexolan 5 cm-pyramidal sound-absorbing foam (Diedorf, Germany) eliminating acoustical reflection. The experiment was controlled by a Hewlett-Packard Intel Core i7 PC computer (Palo Alto, CA, USA) located outside the cabin. The dominant eye’s gaze position was recorded and available online using an EyeLink 1000 Desktop Mounted (SR Research, Osgoode, Ontario, Canada) at a sampling rate of 1 kHz and operated with a 940 nm infrared illuminator. The experimental software controlling the display, the response collection, as well as eye tracking was implemented in Matlab (MathWorks, Natick, MA, USA), using the Psychophysics and EyeLink toolboxes (Brainard, 1997; Cornelissen, Peters, & Palmer, 2002). Stimuli were presented at a viewing distance of 76.5 cm, on a custom-made audiovisual screen (Figure 1A). Auditory distractors were played from 4 loudspeakers (0.75° radius) arranged at the four corners of a virtual square (15° side) centered on the screen midpoint and located at ± 7.5° horizontally and vertically from it. Sounds were played via a multiple channel MOTU sound card (Cambridge, MA, USA), amplified by a PowerPlay Pro Behringer amplifier (Wellich, Germany), and digitized at 48 kHz. Visual distractors were presented via 4 green light emitting diodes (LEDs; 0.15°-radius). Fixation and saccade targets were presented via 5 other red LEDs (0.15°-radius). All LEDs were controlled at a rate of 1 kHz by an Arduino Due electronic board (Turin, Italy). The visual distractors (4 green LEDs) were mounted on top of the 4 speakers. The fixation and saccade targets were located at the center of the screen, at a distance of 15° from the screen center at the four cardinal locations (right, up, left and down). A spandex screen (transparent to sound and light) covered the setup, ensuring that participants remained unaware of stimulus locations before their onset. A custom-made calibration was implemented using the 5 red and 4 green LEDs presented in random sequences. Instructions were recorded and played to participants during training blocks and repeated before each experimental session.

### Procedure

The study was composed of 3 different types of blocks in which we presented visual, auditory or audiovisual distractors, respectively. All blocks were run in a random order across all participants and completed in 3 to 4 experimental sessions (on different days) of about 60-90 minutes each (including breaks). All participants, except one author, initially completed 3 training blocks in which they were familiarized with the task (3 short blocks, one for each distractor type, ~5 minutes each) followed by 12 blocks of the main experiment (3 visual, 3 auditory and 3 audiovisual distractor blocks, ~25 minutes each).

Each trial began with participants fixating a fixation target (red LED) located randomly 15° to the right or left of the screen center (Figure 1B). When the participant’s gaze was detected within a 3.5°-radius virtual circle centered on the fixation target for 200 ms, the trial began with an initial fixation period (varying randomly between 1000 and 1300 ms in steps of 50 ms) followed by the simultaneous presentation of two saccade targets for a duration of 1000 ms (while the fixation target stayed on). The first saccade target always appeared in the center of the screen while the second could randomly appear above or below the screen center. Participants were instructed to make two sequential saccades, the first one toward the screen center, and the second from the screen center to the saccade target. Consequently, participants randomly executed 1 out of 4 possible double-step saccade sequences: left-up, left-down, right-up or right-down saccades. Each trial ended with a 500 ms inter-trial period during which no stimulus was presented.

In 3/4 of the trials, a distractor was presented clockwise or counterclockwise relative to the second saccade target position (e.g. in a right-up saccade trial, a clockwise distractor was presented in the upright quadrant). The onset of the distractor stimulus occurred randomly at a time between 100 ms before and 300 ms after (in steps of 1 ms) the appearance of the saccade targets. This ensured that the distractors reliably occurred either before the first saccade (1st saccade mean latency with visual: 172.02 ± 6.18 ms, auditory: 160.50 ± 6.64 ms, audiovisual: 163.88 ± 5.98 ms or without distractors: 179.44 ± 4.64 ms) or in between the two saccades (2nd saccade mean latency with visual: 497.28 ± 16.21 ms, auditory: 462.92 ± 20.33 ms, audiovisual: 473.86 ± 19.52 ms or without distractor: 480.99 ± 18.84 ms). The distractor within a block remained constant as either a visual, auditory or audiovisual stimulus. A visual distractor was a 50 ms illumination of a green LED, an auditory distractor was a 50 ms broadband Gaussian white noise (with 5 ms raised-cosine onset and offset ramp) and an audiovisual distractor consisted of synchronized visual and auditory distractors originating from the same spatial position. The sound level and frequencies of the speakers were adjusted to be identical (75 dBA SPL) based on records made with a Behringer microphone (Wellich, Germany) placed at the head position. In 1/4 of the trials, no distractor was presented to avoid any difference in saccade preparation. These distractor absent trials were randomly interleaved with distractor present trials. Distractor absent trials were used to normalize the saccade trajectory and thereby account for individual saccade trajectory idiosyncrasies (see *data analysis*).

Participants were instructed to execute the saccades accurately and to avoid looking to the distractor location. To reduce the frequency of saccades made directly from the fixation to the second saccade target (diagonal saccades), we instructed participants to make the requested double-step saccade sequence without strong time pressure. However, a trial was aborted and subjects were given auditory feedback if they took more than 400/700 ms to initiate their first/second saccade relative to the onset of the saccade targets. Each participant completed between 3691 and 3887 trials. Correct fixation as well as correct saccade landing within a 3.5° radius virtual circle centered on the first and second saccade target were monitored online. Trials with fixation breaks or incorrect saccades (inaccurate or too slow) were discarded and repeated at the end of a block in a randomized order (participants repeated between 91 and 287 trials). In total we included 26293 trials (91.30% of the online selected trials, 87.02% of all trials played) in the data analysis.

### Data pre-processing

We first scanned the recorded eye-position data offline and detected saccades based on their velocity distribution (Engbert & Mergenthaler, 2006), using a moving average over twenty subsequent eye position samples. Saccade onset was detected when the velocity exceeded the median of the moving average by 3 SDs for at least 20 ms. We included trials if a correct fixation was maintained within a 3.5° radius centered on the fixation target before the onset of the saccade targets, if a first correct saccade started at the fixation target and landed within a 3.5° radius centered on the first saccade target, if a second correct saccade started at the first saccade target and landed within a 3.5° radius centered on the second saccade target, and if no blink occurred during the trial.

### Data analysis

We first determined the eye position coordinates of the second saccades for each correct trial. These coordinates were then rotated as to direct all saccades upward. Subsequently, we determined the mean eye position on the main direction axis (i.e. mean vertical coordinates across horizontal coordinates) for each second saccade in order to end up with only monotonic eye sample values in the saccade direction. Next, we split data as a function of the direction of the saccade sequence and, for each of these groups, subtracted the mean coordinates of the specific saccade sequence from the mean coordinates of the distractor-absent saccade sequence. This normalization ensured that any deviation of the saccade trajectory in response to a distractor was not due to the individual idiosyncrasy of the saccade trajectory of a participant. It also allowed us to average saccades from different double-step saccade sequences together more accurately. With these normalized coordinates we then determined the saccade curvature angle (Belopolsky & Van der Stigchel, 2013; Jonikaitis & Belopolsky, 2014) for each trial, that is, the median of the angular deviations of each sample point from a straight line connecting the starting and ending point of the saccade. Finally, raw coordinates of the main saccade axis direction were inverted for trials in which distractors were played counterclockwise relatively to the second saccade direction. This way, positive and negative values represent coordinates and curvature angles that were directed either toward or away from the distractor’s head-centered position, respectively. To study the effects of distractors presented before the first saccade, we included trials in which the distractor ended in the last 150 ms preceding the first saccade (pre-saccadic distractor trials, Figure 1C). To study the effects of distractors after the first saccade we included trials in which the distractor started after the first saccade offset and ended at least 100 ms before the second saccade onset (inter-saccadic distractor trials, Figure 1C). Note that the respectively excluded, late occurring distractors (within the last 100 ms prior to the onset of the second saccade) are not expected to be registered early enough by the oculomotor system when processing the second saccade. These time windows were determined following an earlier study on the time course of similar effects (Jonikaitis & Belopolsky, 2014). Per participant, we analyzed 118.38 ± 8.05, 151.12 ± 4.79 and 140.75 ± 4.32 inter-hemifield pre-saccadic, 119.25 ± 6.74, 145.88 ± 4.39 and 138.88 ± 5.62 intra-hemifield pre-saccadic, and 155.00 ± 22.72, 127.75 ± 26.82 and 136.38 ± 26.18 inter-saccadic visual, auditory and audiovisual distractor trials, respectively, and 853.12 ± 21.70 trials without distractor.

Statistical comparisons of normalized saccade curvature angles were based on drawing (with replacement) 10000 bootstrap samples from the original values and computing 10000 means, respectively. By using the bootstrap method, i.e. resampling with replacement from the collected sample, we can form a fair estimate of the parent population. We determined statistical significance by deriving two-tailed *p* values for the comparison of the bootstrapped distributions obtained in a given distractor-present condition to the distributions observed without a distractor. Finally, to compare performance between two distractor-present conditions, we subtracted the bootstrapped values of the first condition from the second and derived two-tailed p-values from the distribution of these differences. We reported uncorrected p-values, as all statistical comparisons were planned and conducted between independent sets of data points.

All analysis codes are available online: https://github.com/mszinte/curvature_av.

## Authors contributions

M.S., D.A.-M., D.J., and H.D designed the experiment. M.S. performed the experiments and analyzed the data. M.S., D.A.-M., D.J., L.W. and H.D discussed the results and wrote the paper.

## Acknowledgments

We are grateful to the members of the Deubel laboratory in Munich for helpful comments and discussions and to Elodie Parison, Alice and Clémence Szinte for their support.

## References

Alais, D., & Burr, D. (2004). The ventriloquist effect results from near-optimal bimodal integration. Current Biology, 14(3), 257–262. http://doi.org/10.1016/j.cub.2004.01.029

Andersen, R. A., Essick, G. K., & Siegel, R. M. (1985). Encoding of spatial location by posterior parietal neurons. Science, 230(4724), 456–458.

Arnott, S. R., & Alain, C. (2011). The auditory dorsal pathway: Orienting vision. Neuroscience & Biobehavioral Reviews, 35(10), 2162–2173. http://doi.org/10.1016/j.neubiorev.2011.04.005

Badde, S., & Heed, T. (2016). Towards explaining spatial touch perception: Weighted integration of multiple location codes. Cognitive Neuropsychology, 33(1-2), 26–47. http://doi.org/10.1080/02643294.2016.1168791

Belopolsky, A. V., & Van der Stigchel, S. (2013). Saccades curve away from previously inhibited locations: evidence for the role of priming in oculomotor competition. Journal of Neurophysiology, 110(10), 2370–2377. http://doi.org/10.1152/jn.00293.2013

Brainard, D. H. (1997). The Psychophysics Toolbox. Spatial Vision, 10(4), 433–436. http://doi.org/10.1163/156856897X00357

Bulkin, D. A., & Groh, J. M. (2012). Distribution of eye position information in the monkey inferior colliculus. Journal of Neurophysiology, 107(3), 785–795. http://doi.org/10.1152/jn.00662.2011

Cavanagh, P., Hunt, A. R., Afraz, A., & Rolfs, M. (2010). Visual stability based on remapping of attention pointers. Trends in Cognitive Sciences, 14(4), 147–153. http://doi.org/10.1016/j.tics.2010.01.007

Chen, X., DeAngelis, G. C., & Angelaki, D. E. (2013). Diverse spatial reference frames of vestibular signals in parietal cortex. Neuron, 80(5), 1310–1321. http://doi.org/10.1016/j.neuron.2013.09.006

Colby, C. L., Duhamel, J. R., & Goldberg, M. E. (1996). Visual, presaccadic, and cognitive activation of single neurons in monkey lateral intraparietal area. Journal of Neurophysiology, 76(5), 2841–2852.

Collins, T., Heed, T., & Röder, B. (2010). Eye-movement-driven changes in the perception of auditory space. Attention, Perception, & Psychophysics, 72(3), 736–746. http://doi.org/10.3758/APP.72.3.736

Cornelissen, F. W., Peters, E. M., & Palmer, J. (2002). The Eyelink Toolbox: Eye tracking with MATLAB and the Psychophysics Toolbox. Behavior Research Methods, Instruments, & Computers, 34(4), 613–617. http://doi.org/10.3758/BF03195489

Doyle, M., & Walker, R. (2002). Multisensory interactions in saccade target selection: Curved saccade trajectories. Experimental Brain Research, 142(1), 116–130. http://doi.org/10.1007/s00221-001-0919-2

Duhamel, J. R., Colby, C. L., & Goldberg, M. E. (1992). The Updating of the Representation of Visual Space in Parietal Cortex by Intended Eye-Movements. Science, 255(5040), 90–92.

Engbert, R., & Mergenthaler, K. (2006). Microsaccades are triggered by low retinal image slip. Proceedings of the National Academy of Sciences of the United States of America, 103(18), 7192–7197. http://doi.org/10.1073/pnas.0509557103

Frens, M. A., & Van Opstal, A. J. (1995). A quantitative study of auditory-evoked saccadic eye movements in two dimensions. Experimental Brain Research, 107(1), 103–117.

Frens, M. A., Van Opstal, A. J., & Van Der Willigen, R. F. (1995). Spatial and temporal factors determine auditory-visual interactions in human saccadic eye movements, 57(6), 802–816. http://doi.org/10.3758/BF03206796

Godfroy-Cooper, M., Sandor, P. M. B., Miller, J. D., & Welch, R. B. (2015). The interaction of vision and audition in two-dimensional space. Frontiers in Neuroscience, 9, 70. http://doi.org/10.3389/fnins.2015.00311

Goossens, H. H., & Van Opstal, A. J. (1999). Influence of head position on the spatial representation of acoustic targets. Journal of Neurophysiology, 81(6), 2720–2736.

Gottlieb, J. P., Kusunoki, M., & Goldberg, M. E. (1998). The representation of visual salience in monkey parietal cortex. Nature, 391(6666), 481–484.

Groh, J. M., Trause, A. S., Underhill, A. M., Clark, K. R., & Inati, S. (2001). Eye position influences auditory responses in primate inferior colliculus. Neuron, 29(2), 509–518.

Grunewald, A., Linden, J. F., & Andersen, R. A. (1999). Responses to auditory stimuli in macaque lateral intraparietal area. I. Effects of training. Journal of Neurophysiology, 82(1), 330–342.

Heeman, J., Nijboer, T. C. W., Van der Stoep, N., Theeuwes, J., & Van der Stigchel, S. (2016). Oculomotor interference of bimodal distractors. Vision Research, 123, 46–55.

Henriques, D. Y. P., Klier, E. M., Smith, M. A., Lowy, D., & Crawford, J. D. (1998). Gaze-Centered Remapping of Remembered Visual Space in an Open-Loop Pointing Task. The Journal of Neuroscience, 18(4), 1583–1594.

Jay, M. F., & Sparks, D. L. (1984). Auditory receptive fields in primate superior colliculus shift with changes in eye position. Nature, 309(5966), 345–347.

Jay, M. F., & Sparks, D. L. (1987). Sensorimotor integration in the primate superior colliculus. II. Coordinates of auditory signals. Journal of Neurophysiology, 57(1), 35–55.

Jonikaitis, D., & Belopolsky, A. V. (2014). Target-distractor competition in the oculomotor system is spatiotopic. The Journal of Neuroscience, 34(19), 6687–6691. http://doi.org/10.1523/JNEUROSCI.4453-13.2014

King, A. J. (2004). The superior colliculus. Current Biology, 14(9), R335–R338. http://doi.org/10.1016/j.cub.2004.04.018

King, A. J., Hutchings, M. E., Moore, D. R., & Blakemore, C. (1988). Developmental plasticity in the visual and auditory representations in the mammalian superior colliculus. Nature, 332(6159), 73–76. http://doi.org/10.1038/332073a0

King, A. J., Schnupp, J. W. H., & Doubell, T. P. (2001). The shape of ears to come: dynamic coding of auditory space. Trends in Cognitive Sciences, 5(6), 261–270.

Klier, E. M., Wang, H., & Crawford, J. D. (2001). The superior colliculus encodes gaze commands in retinal coordinates, 4(6), 627–632. http://doi.org/10.1038/88450

Kruijne, W., Van der Stigchel, S., & Meeter, M. (2014). A model of curved saccade trajectories: spike rate adaptation in the brainstem as the cause of deviation away. Brain and Cognition, 85, 259–270. http://doi.org/10.1016/j.bandc.2014.01.005

Krüger, H. M., Collins, T., Englitz, B., & Cavanagh, P. (2016). Saccades create similar mislocalizations in visual and auditory space. Journal of Neurophysiology, 115(4), 2237–2245. http://doi.org/10.1152/jn.00853.2014

Kusunoki, M., & Goldberg, M. E. (2003). The Time Course of Perisaccadic Receptive Field Shifts in the Lateral Intraparietal Area of the Monkey. Journal of Neurophysiology, 89(3), 1519–1527.

Lee, J., & Groh, J. M. (2012). Auditory signals evolve from hybrid- to eye-centered coordinates in the primate superior colliculus. Journal of Neurophysiology, 108(1), 227–242. http://doi.org/10.1152/jn.00706.2011

Linden, J. F., Grunewald, A., & Andersen, R. A. (1999). Responses to auditory stimuli in macaque lateral intraparietal area. II. Behavioral modulation. Journal of Neurophysiology, 82(1), 343–358.

Maddox, R. K., Pospisil, D. A., Stecker, G. C., & Lee, A. K. C. (2014). Directing Eye Gaze Enhances Auditory Spatial Cue Discrimination. Current Biology, 24(7), 748–752.

Melcher, D., & Colby, C. L. (2008). Trans-saccadic perception. Trends in Cognitive Sciences, 12(12), 466–473. http://doi.org/10.1016/j.tics.2008.09.003

Nakamura, K., & Colby, C. L. (2002). Updating of the visual representation in monkey striate and extrastriate cortex during saccades. Proceedings of the National Academy of Sciences, 99(6), 4026–4031. http://doi.org/10.1073/pnas.052379899

Pavani, F., Husain, M., & Driver, J. (2008). Eye-movements intervening between two successive sounds disrupt comparisons of auditory location. Experimental Brain Research, 189(4), 435–449. http://doi.org/10.1007/s00221-008-1440-7

Populin, L. C., Tollin, D. J., & Yin, T. C. T. (2004). Effect of eye position on saccades and neuronal responses to acoustic stimuli in the superior colliculus of the behaving cat. Journal of Neurophysiology, 92(4), 2151–2167. http://doi.org/10.1152/jn.00453.2004

Robinson, D. A. (1972). Eye movements evoked by collicular stimulation in the alert monkey. Vision Research, 12(11), 1795–1808. http://doi.org/10.1016/0042-6989(72)90070-3

Robinson, D. A., & Fuchs, A. F. (1969). Eye movements evoked by stimulation of frontal eye fields. Journal of Neurophysiology, 32(5), 637–648.

Rolfs, M., & Szinte, M. (2016). Remapping Attention Pointers: Linking Physiology and Behavior. Trends in Cognitive Sciences, 20(6), 399–401. http://doi.org/10.1016/j.tics.2016.04.003

Russo, G. S., & Bruce, C. J. (1994). Frontal eye field activity preceding aurally guided saccades. Journal of Neurophysiology, 71(3), 1250–1253.

Sheliga, B. M., Riggio, L., & Rizzolatti, G. (1994). Orienting of attention and eye movements. Experimental Brain Research, 98(3), 507–522. http://doi.org/10.1007/BF00233988

Sommer, M. A., & Wurtz, R. H. (2006). Influence of the thalamus on spatial visual processing in frontal cortex. Nature, 444(7117), 374–377. http://doi.org/10.1038/nature05279

Sparks, D. L., & Nelson, I. S. (1987). Sensory and motor maps in the mammalian superior colliculus. Trends in Neurosciences, 10(8), 312–317. http://doi.org/10.1016/0166-2236(87)90085-3

Stricanne, B., Andersen, R. A., & Mazzoni, P. (1996). Eye-centered, head-centered, and intermediate coding of remembered sound locations in area LIP. Journal of Neurophysiology, 76(3), 2071–2076.

Szinte, M., Jonikaitis, D., Rangelov, D., & Deubel, H. (2018). Pre-saccadic remapping relies on dynamics of spatial attention. eLife, 7, e37598.

Theeuwes, J., Olivers, C. N. L., & Chizk, C. L. (2005). Remembering a location makes the eyes curve away. Psychological Science, 16(3), 196–199. http://doi.org/10.1111/j.0956-7976.2005.00803.x

Van der Stigchel, S., Meeter, M., & Theeuwes, J. (2006). Eye movement trajectories and what they tell us. Neuroscience & Biobehavioral Reviews, 30(5), 666–679. http://doi.org/10.1016/j.neubiorev.2005.12.001

Van Grootel, T. J., Van Wanrooij, M. M., & Van Opstal, A. J. (2011). Influence of static eye and head position on tone-evoked gaze shifts. The Journal of Neuroscience, 31(48), 17496–17504. http://doi.org/10.1523/JNEUROSCI.5030-10.2011

Vliegen, J., Van Grootel, T. J., & Van Opstal, A. J. (2004). Dynamic sound localization during rapid eye-head gaze shifts. The Journal of Neuroscience, 24(42), 9291–9302. http://doi.org/10.1523/JNEUROSCI.2671-04.2004

Walker, M. F., Fitzgibbon, E. J., & Goldberg, M. E. (1995). Neurons in the monkey superior colliculus predict the visual result of impending saccadic eye movements. Journal of Neurophysiology, 73(5), 1988–2003.

Wurtz, R. H. (2008). Neuronal mechanisms of visual stability. Vision Research, 48(20), 2070–2089. http://doi.org/10.1016/j.visres.2008.03.021

